# Zerumbone From *Zingiber zerumbet* (L.) Smith: A Potential Prophylactic and Therapeutic Agent Against The Cariogenic Bacterium *Streptococcus mutans*

**DOI:** 10.1101/187906

**Authors:** Thiago Moreira da Silva, Carlos Danniel Pinheiro, Patricia Puccinelli Orlandi, Carlos Cleomir Pinheiro, Gemilson Soares Pontes

**Author notes:** **Corresponding author** PhD. Gemilson Soares Pontes, National Institute of Amazonian Research, Society Environment and Health Department, Laboratory of Virology, Manaus, Amazonas, Brazil.

## Abstract

**Background:** Essential oil obtained from rhizomes of the *Zingiber zerumbet* (L.) Smith (popularly known in Brazil as bitter ginger) is mainly constituted by the biomolecule zerumbone, which exhibit untapped antimicrobial potential. The aim of this study was to investigate the antimicrobial activity of the zerumbone from bitter ginger rhizomes against the cariogenic agent *Streptococcus mutans*.

**Methods:** Firstly, the essential oil from rhizomes of *Zingiber zerumbet* (L.) Smith extracted by hydrodistillation was submitted to purification and recrystallization process to obtain the zerumbone compound. The purity of zerumbone was determined through high-performance liquid chromatography analysis. Different concentrations of zerumbone were tested against the standard strain *S. mutans* (ATCC 35668) by using microdilution method. The speed of cidal activity was determined through a time kill-curve assay. The biological cytotoxicity activity of zerumbone was assessed using Vero Cell line through MTT assay.

**Results:** The zerumbone showed a minimum inhibitory concentration (MIC) of ≥ 250μg/ mL and a minimum bactericidal concentration (MBC) of ≥ 500μg/mL against *S. mutans*. After six hours of bacteria-zerumbone interaction, all concentrations tested starts to kill the bacteria and all bacteria were killed between 48 and 72 hours period at the concentration of 500 μg/ mL (99,99% of bacteria were killed in comparison with original inoculum). In addition, zerumbone showed no cytotoxicity activity on mammalian continuous cells line.

**Conclusions:** These results draw attention to the great potential of zerumbone as antimicrobial agent against *S. mutans* infection, indicating its possible use in the phyto-pharmaceutical formulations as new approach to prevent and treat tooth decay disease.

Abbreviations

MIC: Minimum Inhibitory Concentration
MBC: Mininum Bactericidal Concentration
CFU: Colony forming units
BHI: Brain Heart Infusion
RPM: rotation per minute
OD: optical density

## Background

Several medicinal plants contain in their biochemical constitution many compounds with antibacterial action that remains unknown. *Zingiber zerumbet* (L.) Smith, a rhizomatous herbaceous specie, that belongs to the family *Zingiberaceae,* is a native plant from Southeast Asia with potential antimicrobial activity not fully comprehended [1]. In Brazil, this specie is usually identified as bitter ginger and it is easily found in large amount at Amazonas state, where it is well adapted to local climate conditions.

Many types of plant essential oils have been reported as a source of biomolecules with promising antimicrobial activities [2,3].Essential oils from *Zingiber zerumbet* rhizomes have great potential pharmacological activities, including antimicrobial, anti-inflammatory, chemo-preventive, antinociceptive, antiulcer, antioxidant, antipyretic and analgesic, as previously described [4–8].However, its antimicrobial activity spectrum remains to be determined.

The major bioactive molecule found in the essential oil of *Zingiber zerumbet* rhizomes is the zerumbone (figure 1), a monocyclic sesquiterpene compound (2,6,10-cy-cloundecatrien-1-one, 2,6,9,9-tetramethyl-,(E,E,E)-) [9]. Zerumbone has been linked to a broad of biological activities, including antibacterial action [10,11]. Previous reports demonstrated the antimicrobial activity of the zerumbone against gram negative bacteria, such as *Escherichia coli* and *Helicobacter pilory*, and gram-positive bacteria, such as *Staphylococcus epidermidis* and *aureus*, showing more antibacterial effectiveness on gram positive microorganisms [8,12].Besides that, more studies are needed to determine the potential of the zerumbone as antibacterial agent against gram positive microorganism, like the *Streptococcus mutans,* the main agent of tooth decay, the oral infectious disease most prevalent in the world that affects over 90% of school-aged children and about 100% of the world population [13].

**Figure 1:**
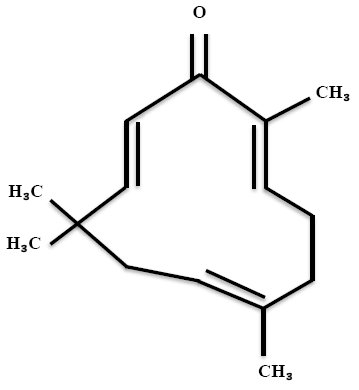
Chemical structure of zerumbone.

Despite the great diversity of bacterial species in the oral cavity, few are able to cause tooth decay (cariogenic bacteria) and the *S. mutans* has been implicated as the major causative agent of this oral infectious disease [14].The cariogenic potential displayed for this bacterium is due to its ability to produce acid (acidogenic) from dietary carbohydrate, capacity to survive in low-pH environments (aciduric) and, especially, due to its great ability to adhere onto the tooth surfaces, which makes the *S. mutans* responsible for the initial formation of dental plaque [15].

Some chemicals agents available such as chlorexidin and others phenolic compounds are usually used to prevent the tooth decay, but the long-term use may result in side effects as loss of taste, metallic taste in mouth, dental pigmentation, diarrhea and an oral burning sensation [16]. On this context, biomolecules isolated from plants have been suggested as an alternative therapeutic over synthetic chemical agents for prevention of tooth decay, because of their few or no side effects [17]. Thus, the main goal of this study was to investigate the antimicrobial activity of the zerumbone obtained from rhizomes of *Zingiber zerumbet* (L.) Smith against *S. mutans*, the main etiological agent of tooth decay.

## Materials and Methods

### Acquisition of *Zingiber zerumbet* rhizomes

The rizhomes of the *Z. zerumbet* were collected in a rural area surrounding the city of Manaus/AM, located at BR-174, points P01 to P02, latitude 24132, 03789 S and longitude 600931,40854 W, according to geographic coordination. After that, an exsiccate was sent to the herbarium of the National Institute of Amazonian Research (INPA) for proper identification and comparison with the exsiccate previously identified by Prof. Dr. Paul Maas (Department of Plant Ecology and Evolutionary Biology; Herbarium University of Utrecht), which is deposited in the herbarium under N°. 186913.

### Essential oil extraction

The extraction of the essential oil (EO) was carried out in the Thematic Laboratory of Chemistry and Natural Products at INPA. The EO was obtained through the hydrodistillation of rhizomes. Briefly, after the identification, cleaning and disinfection, the material was crushed and dried at room temperature. Clevenger apparatus adapted with a round-bottom flask of 2 liters volume was used for distillation of OE from the crushed material diluted in distilled water in a proportion 1:4. The extraction was done during 6 hours starting at the boiling point. Afterward, the EO was collected from the condenser and stored in amber flasks at room temperature. All system was protected from light using aluminum foil. The OE yield was estimated by calculating the ratio between the oil mass and the feed mass.

### Gas chromatography–mass spectrometry (GC–MS) analysis

The analysis of the essential oil composition was carried out using Hewlett Packard HP/série 6890 GC SITEM PLUS gas chromatographer (6511 Bunker Lake Blvd. Ramsey, Minnesota, 55303 USA) with an analytical HP-5MS 5% phenylmethylsiloxane capillary column (30 m x0.32 mm i.d, film thickness 0,25 μm). It was operated an electron ionization system with ionization energy of 70 V. The analyses were conducted using helium and nitrogen gases at 2 mL/min with purity percentage of 99.999%. It was injected 1 μL in splitless mode, in 1:20 ratio of hexannic solution. The oven was programmed at controlled temperature between 60-240° C, raising 3°C/min and kept at 250°C for 10 minutes. Temperature of mass transfer and injection were established at 220° C and 290° C, respectively. The chemical constituents of essential oil analyzed were expressed as relative percentage by peak area normalization. The identification of the OE constituents was done by calculating the retention time obtained in the analyzes of GC-MS, correlating them with the retention times of the n-alkanes (C9-C30). The indices were compared with the data available in the NIST / WILEY library [18]. The zerumbone identification was done as previously described [19].

### High-performance liquid Chromatography (HPLC)

After purification and recrystallization of the EO using our patented method (n^0^ PI-0505343-9/28/11/2007), the zerumbone purity was estimated through HPLC analysis (Accela High Speed LC, Thermo Scientific®), using a column Hypersil Gold (50 × 2,1 mm) and a mobile phase methanol:water (85:15, v/v) at 1 mL/min. The identification was done by comparing the retention time of the peaks with those standard solutes in HPLC and confirmed by UV-absorption spectrum (∼252 nm).

### Inoculum standardization

The antimicrobial activity of the zerumbone was evaluated against the standard strain of *S. mutans* ATCC 35668 (American Type Culture Collection, Microbiologics Inc., St. Cloud, USA). Firstly, bacterial suspensions were prepared by inoculation of colonies into a tube containing 3 mL Brain Heart Infusion (BHI) broth, followed by incubation at orbital shaking of 150 RPM for 72 hours at 37 °C in anaerobic conditions, as previously described [20]. After incubation, the turbidity was calibrated and adjusted through spectrophotometer analysis to match the 0,5 MacFarland scale (1×10^8^ CFU/mL). Final inoculum of 1 × 10^6^ CFU/ mL was used in the assays.

### Determination of the Antimicrobial Activity

The antimicrobial activity of zerumbone was evaluated by estimation of Minimum Inhibitory Concentration (MIC) and Minimum Bactericidal Concentration (MBC) using microdilution method according to the guidelines of the Clinical & Laboratory Standards Institute (CLSI)[21]. The concentrations of zerumbone tested were 2000; 1000; 500; 250 and 125 μg/mL. After diluting serially the zerumbone in 96 well-plate containing 100 μL BHI broth, 100 μL of BHI broth having bacterial inoculum (1×10^6^CFU/mL) were added into wells resulting in a final volume of 200 μL, followed by incubation at the same conditions mentioned above. Additional wells containing diluting agent of zerumbone (tween 20 10%) was used as a control.

After incubation, the bacterial growth was evaluated by measuring the turbidity in each well through spectrophotometric analysis (600nm). Subsequently, an aliquot of 50 μL of each well was collected and seeded on plates containing BHI agar and incubated for 72 hours at 37 °C in anaerobic conditions. Next, the plates were analyzed for the presence/absence of bacteria in order to estimate the MIC and MBC. All tests were performed in triplicate and repeated three times to verify the reproducibility of results.

### Time-Kill Curve Assay

To determine the speed of cidal activity of the zerumbone, a time kill-curve was performed as previously described, with few modifications[22]. An inoculum of 1×10^6^ CFU/mL was added into tubes having 3 mL of BHI broth or BHI broth treated with zerumbone (250 or 500 μg/mL) or tween 20 10% (control) and then incubated for 72 hours anaerobically with orbital rotation of 150 RPM at 37 °C. The tube containing only bacteria and BHI broth was used to estimate the different phases of growth curve.

An aliquot of 100 μL was removed at 0, 6, 12, 24, 48 and 72 h time intervals to determine the bacterial growth by measuring turbidity through spectrophotometric analysis (600nm). Subsequently, the aliquots were serially diluted in 0.85% of sterile saline solution, seeded in BHI agar and then incubated at the conditions set out above. The viable number of bacterial cells was estimated by counting CFU and multiplying the results by dilution factors. Means of duplicate colony counts were taken. To build the time-kill curve, the log_10_ CFU/mL versus time over 72 h was plotted. The decrease of 99.9% (≥ 3 log_10_) of the total number of CFU/mL in the original inoculum was used to estimate the bactericidal activity. The assays were performed in triplicate and repeated three times to confirm the reproducibility of results.

### Cytotoxicity assay

The cytotoxicity activity of zerumbone was determined using the MTT (3-(4, 5-dimethyl thiazol-2-yl)-2, 5-diphenyl tetrazolium bromide) assay as previously described with few modifications [20]. Summarily, VERO cells line (2 × 10^4^ per well) were cultured into 96-well plate containing 0.2mL of DMEM medium (with 10% FBS, penicillin-streptomycin and amphotericin B) per well, in atmosphere of 5% CO_2_ at 37° C for 24 hours. After formation of sub-confluent monolayer, the cells were treated with zerumbone (25 μg/mL; 50 μg/mL and 100 μg/mL) and incubated again at the same conditions above mentioned for 24 and 48 hours. Sterile PBS was used as positive control and DMSO 100% as negative control. Subsequently, the medium was removed from all wells and 10μL of MTT (5 mg/mL in sterile PBS) diluted in 100 μL of DMEM medium (without phenol red to avoid misinterpretation) was added into the wells and incubated in atmosphere of 5% CO_2_ at 37° C for 4 hours. After that, the MTT was removed and 50μL of MTT lysis buffer were added to each well followed for gently homogenization to dissolve the formazan crystals and incubated again for 10 minutes at the same conditions mentioned earlier. Optical densities of samples were measured using a microplate reader at wavelength of 570 nm. The relative viability of cells was estimated using the following equation: (A570 of treated sample)/(A570 of untreated sample)×100. All tests were done in triplicate.

## Results

### Acquisition of Essential oil and purification of zerumbone

The yield of the essential oil obtained was 5%. The EO showed to be constituted mainly by sesquiterpene zerumbone compound, according to GC-MS analysis (figure 2). A percentage of 87,93% of zerumbone was detected among the nineteen others chemical constituents. The purification and recrystallization processes applied to EO resulted in a zerumbone crystal with 98% of purity (figure 3).These crystals were used in all subsequent tests.

**Figure 2:**
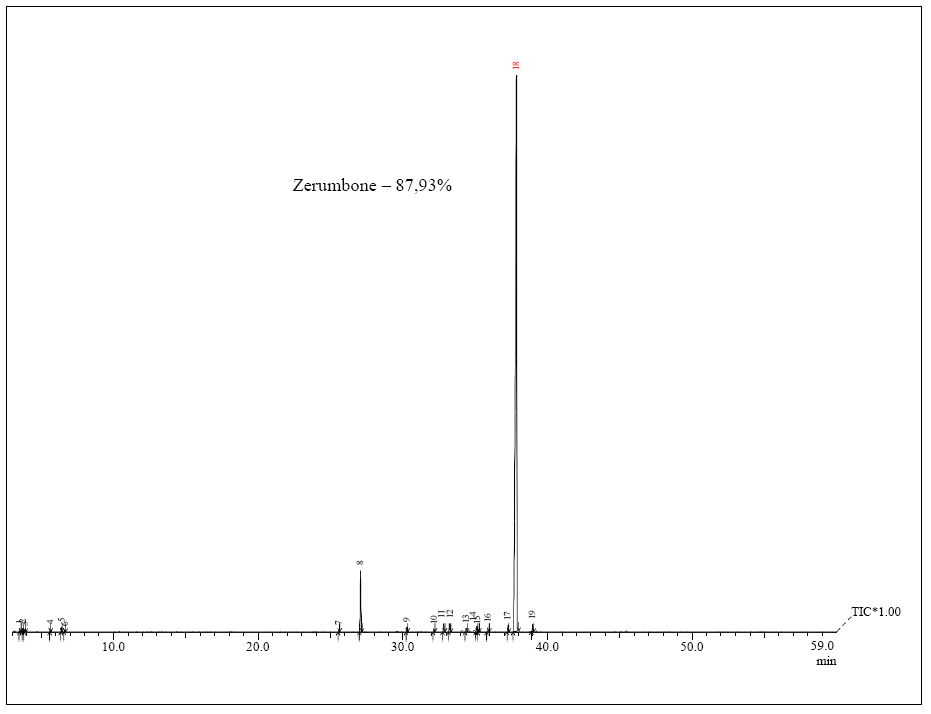
Gas Chromatography-Mass Spectrometry (GC-MS) of Essential oil *Zingiber zerumbet* (L.) Smith. Analysis revealed the presence of 19 compounds. Zerumbone was the major compound (87,93%) found in the essential oil used in this study.* R. time: retention time.

**Figure 3:**
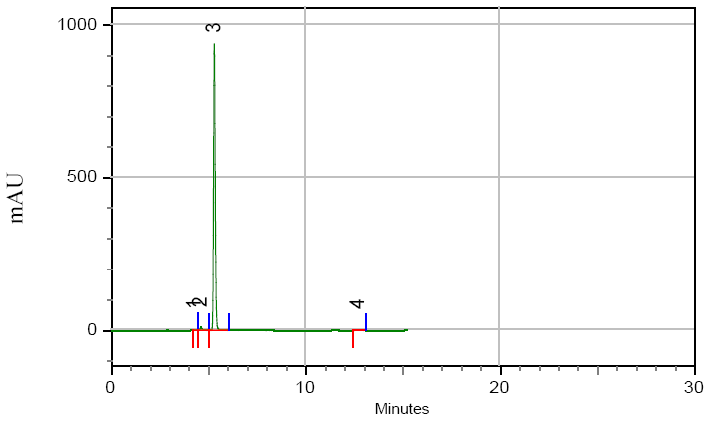
High-performance liquid Chromatography (HPLC) of zerumbone crystals. HLPC elution of zerumbone crystal with retention time of ∼5 minutes confirmed zerumbone purity of 98% (peak 3).

### Antimicrobial activity of zerumbone

The zerumbone demonstrated efficient antimicrobial activity against *S. mutans* ATCC 35668 showing a MIC value of ≥ 250 μg/mL and MBC of ≥ 500 μg/mL (Figure 4). The MIC was determined according to lowest concentration of essential oil that inhibits the bacterial growth, whereas the MBC was assessed based on the concentration that kills all viable bacterial cells (supplementary figure 2).

**Figure 4:**
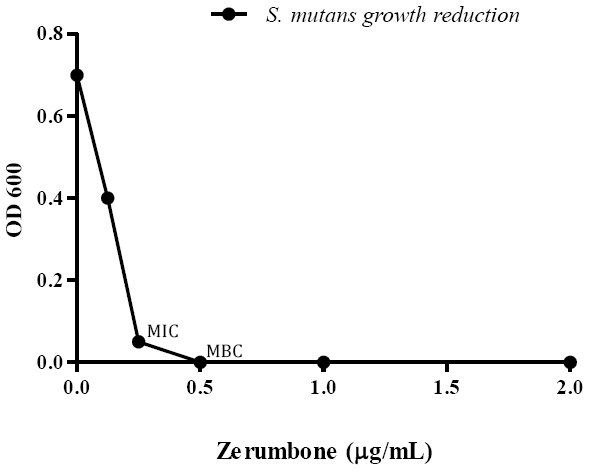
Antimicrobial activity (MIC and MBC) of zerumbone against *S. mutans*. Bacterial inoculum (1×10^6^ CFU/mL) was treated with different concentrations of zerumbone and incubated at 37° C for 48 h in anaerobic conditions. After incubation, the bacterial growth was verified by turbidity measurements using spectrophotometer.The results are representative of three independent experiments performed in triplicate. OD: optical density; MIC: Minimum Inhibitory Concentration; MBC: Minimum bactericidal concentration

### Time-kill Curve

The time-kill curve assay corroborates the antimicrobial activity of the zerumbone. The zerumbone displayed more intense antibacterial activity in the time interval 12-48 hours, corresponding to log phase of *S. mutans* growth curve (supplementary figure 2). Within 12 hours of bacterial exposure to 250 μg/mL and 500 μg/mL of zerumbone concentrations, viable bacterial cells were reduced in 41,77% and 58,34%, respectively, in comparison to the original inoculum (figure 4 A). After 24 hours of exposure, the reduction was more pronounced with 64.29% (250 μg /mL) and 78.58% (500 μg/mL) of bacterial cells death (figure 4 B and C).Finally, the zerumbone showed its maximum action in the interval of 48-72 hours, reducing 96,88% of bacterial colonies at concentration of 250 μg/mL and killing all bacteria at the concentration of 500 μg/mL (Figure 5 A, B and C).

**Figure 5:**
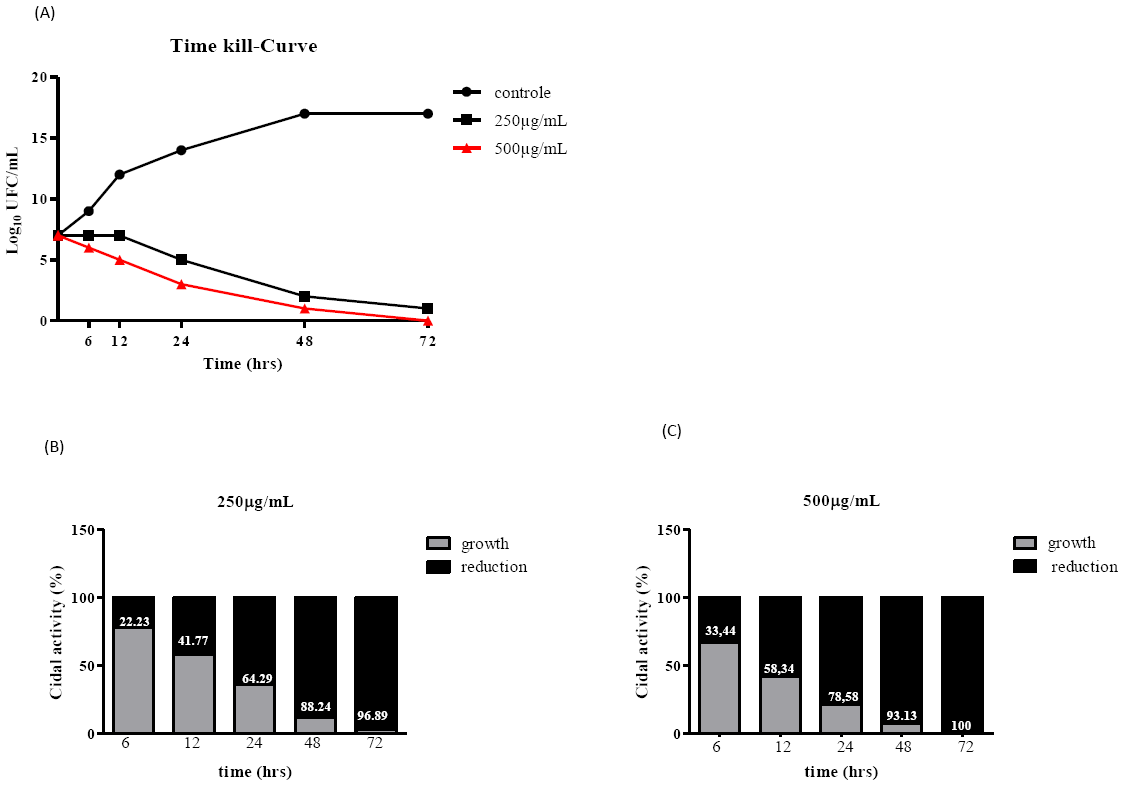
Bacterial‐ time kill curve. Zerumbone was tested for antimicrobial activity on bacterial-kill kinetics. *S. mutans* were grown in BHI broth + tween 20 10% (control) or along with graded concentrations of zerumbone at 37°C for 72h in anaerobic conditions. Samples were collected at different time intervals to estimate bacterial kill kinetics (A) and growth reduction (B and C), by spectrophotometry analysis and CFU counts. Decrease of 2log_10_ at any time point from original Inoculum was considered significant. Results are representative of three independent experiments. Numbers inside of bars at (B) and (C) figures means reduction percent.

### Cytotoxicity activity of zerumbone

The MMT assay demonstrated that the biomolecule zerumbone had no considerable cytotoxicity effect up to 100 μg/mL. At concentrations 25, 50 and 100 μg/mL the percentages of cell viability were 100, 97 and 92%, respectively, after 24 hours treatment (figure 6). The cell viability slightly changed at concentration 50 μg/mL (95%) and 100 μg/mL (85%) after 48 hours treatment. The results clearly show that the zerumbone may have no cytotoxic effect on normal mammalian cells.

**Figure 6:**
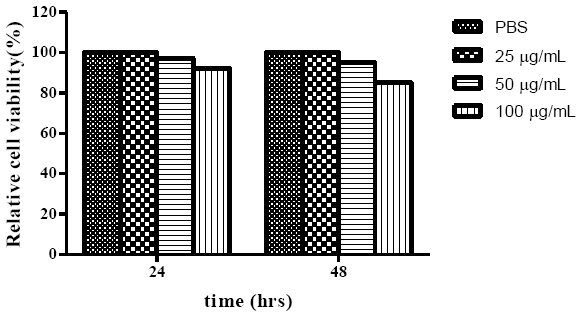
Cytotoxicity of zerumbone against Vero cell line using MTT assay.

## Discussion

Tooth decay disease is a growing global concern that threatens human health and safety [23]. Emerging resistance to conventional antibiotics associated to inconvenient side effects caused by commercial antibiotics usually result in treatment leaving [24]. In this regard, the development of alternative antibiotic therapies based on bioactive molecules from medicinal plants is urgent and crucial to provide effective prevention and treatment for tooth-infection-causing bacteria.

This study demonstrated strong antimicrobial activity of zerumbone against the cariogenic agent *S. mutans* (MIC> 250 μg/ mL) according to the following classification of antimicrobial action: strong = 50<MIC<500 μg/mL moderate = MIC 600<MIC<1500 μg/mL and; weak = MIC>1500 μg/mL [25,26]. Thus, our data bring to the light a new possibility of prophylactic and therapeutic strategies for *S. mutans* infection, especially in case of tooth decay, a serious public health issue [27].

As a matter of fact, few reports have shown natural biomolecules antimicrobial activity against the *S. mutans* and the majority demonstrated weak antibacterial action based on the classification set out above [28–30].A recent study tested around two thousand plant extracts from Amazon region against *S. mutans* (ATCC 25175) and, from this number, seventeen displayed antimicrobial activity, but only the extract obtained from *Ipomoea alba* L. sp (*Convolvulaceae*) showed antimicrobial activity considered strong (MBC> 160μg/mL) [31].Takarada et al. [32] and Aguiar et al. [33] evaluated the anti-*S. mutans* activity of essential oils from the following plants: *Romarinus officinalis* L., *Melaleuca alternifólia*, *Lavandula officinalis*, *Leptosperfum scoparium*, *Eucalyptus radiate*, *Ageratum conyzoides*, *Artemisia camphorata* Vill., *Bidens sulphurea*, *Foeniculum vulgare* Mill., *Lippia alba*, *Ocimum gratissimum* L., *Pelargonium graveolens*, *Syzygium aromaticum* and *Tagetes erecta* L. The results showed MICs ranging from 500 to10,000 μg/mL, activities considered weak compared to our data. Therefore, these studies corroborate the strong bioactive potential of the zerumbone as anticariogenic agent, which could be a good substrate to be used in prophylactic and therapeutic formulations against tooth decay.

Indeed, the zerumbone has been tested against a range of microorganisms showing good antibacterial activity mainly against gram positive bacteria [34]. But, up to now, there is no report describing the antimicrobial activity of the zerumbone or its analogs against *S. mutans*. Hasan et al. [35] demonstrated that extracts from *Zingiber officinale*, specie that belongs to same genus and family of bitter ginger, has anti-*S. mutans* action with MIC value of 256 μg/mL, effect similar to the one observed for this study. However, differently from our study, no compound was isolated or purified to determine which biomolecules were responsible for the antimicrobial activity observed. Generally, the essential oil obtained from *Zingiber zerumbet* shows in its constitution a range of zerumbone concentration starting from 12 to 73%[9]. Our results demonstrated that the essential oil obtained in this study showed 87,93% of zerumbone, which was raised to 98% after purification and recrystallization process using a method patented for our group (n^0^ PI-0505343-9/28/11/2007). This purity degree usually is greater than the ones described elsewhere, which allow us to better characterize the zerumbone antimicrobial action [10].

As the zerumbone exhibited efficient antimicrobial activity, we next evaluated the speed of cidal activity. The effectiveness observed was concentration and time-dependent, although both concentrations tested showed similar results. The intensity of zerumbone antimicrobial activity was more pronounced during the logarithmic growth phase. After 24 hours of bacterial exposure to 250 and 500 μg/mL zerumbone concentrations, viable bacterial cells were reduced 64.29% and 78.58%, respectively. According to Jones et al. [36] when an antimicrobial agent reduces the bacterial colonies around 70% within 24 hour time period, it can be considered as a strong candidate for treating bacterial infections. However, the maximum cidal activity of zerumbone was reached at stationary phase, suggesting the possibility of two mechanism of zerumbone activity, different from the most antibiotic agents that require cell division or active metabolism for the drug’s killing activity [37,38]. Thus, an antibiotic agent that still shows bactericidal activity under growth-limited conditions may be very advantageous, since the biofilm-associated microorganisms, such as *S. mutans*, may modulate gene expression to enhance endurance under periods of nutrient limitation to survive in the stationary phase [39]. Nevertheless, more studies are needed to elucidate the zerumbone antibacterial mechanism of action.

Extracts derived from *Dodonaea viscosa* plant killed 100% of *S. mutans* colonies within 24 h exposition [40]. However, the concentration required to kill all bacteria was 12.500 mg/mL, value much higher than the one observed in this study. Nonetheless, it is important to emphasize that comparison of time-kill curve among different bacteria strains should be done carefully once the microbial growth logarithmical phase directly influences the action of antibacterial agent [41].

Beyond the determination of the optimum antibacterial concentration, it is also important guarantee the cytotoxicity safety of the bioactive substances against normal mammalian cells, if the ultimate goal is to use it as a main raw material to produce new drugs. Usually, finding the balance between effectiveness and safety of a biomolecule it is very hard mission [42,43].Our findings indicated that the zerumbone is not cytotoxic to normal mammalian cells, since the exposure of VERO cells to different concentrations of zerumbone cause no considerable toxic effects. Even after 48 hours of zerumbone treatment, no substantial toxic effect was observed, other than a low cytotoxicity evidenced at 100μg/mL concentration (reduction of 15% of cell proliferation). However, it is imperative to consider that the susceptibility of mammalian cells tend to be greater in the MTT test than *in vivo* situation, because of the direct exposure of the cells to the biomolecules without any type of variants happening *in vivo*, such as route of administration and topical absorption, which may influence or even though decrease the cytotoxicity effect demonstrated *in vitro* [44].

Altogether, this study demonstrated the zerumbone great antimicrobial activity against the *S. mutans*, showing that its toxic activity is selectively targeted to the bacterium, once the zerumbone did not display cytotoxic effect against the normal mammalian cells. However, further studies using *in vitro* and *in vivo* models are needed to better determine the zerumbone effectiveness and safety as antimicrobial agent in the context of prophylaxis and treatment of cariogenic infections.

## Conclusions

In summary, this study suggests that the zerumbone represents an excellent bioactive substance to be explored by phytopharmaceutical industry in the drug formulations, to prevent and treat cariogenic infections, because this biomolecule showed an outstanding antimicrobial activity against the main etiological agent of tooth decay, *S. mutans.* Although the cytotoxicity safety of zerumbone in mammalian cells and its antibacterial action at both log and stationary phases support its potential use as antibacterial agent against *S. mutans* infection, additional studies are necessary to better characterize the zerumbone mechanism of action, efficacy and safety in the scenario of cariogenic disease.

## Acknowledgements

We would like to express our sincere gratitude to the Thematic laboratory of Chemistry and Natural Products of National Institute of Amazonian Research to all support provided through the execution of this research.

## Availability of data and materials

All the details of data and materials of this study are included in the manuscript.

## Authors’ contributions

TMS, CCP, CDP carried most of the experiments; TMS, GSP, PPO wrote the manuscript. GSP, TMS and CCP design the project structure. All authors read and approved the final manuscript.

## Ethics approval and consent to participate

Not applicable

## Consent for publication

All co-authors have consented for the publication of this manuscript.

## 10. Figures legends

**Supplementary figure 1.**
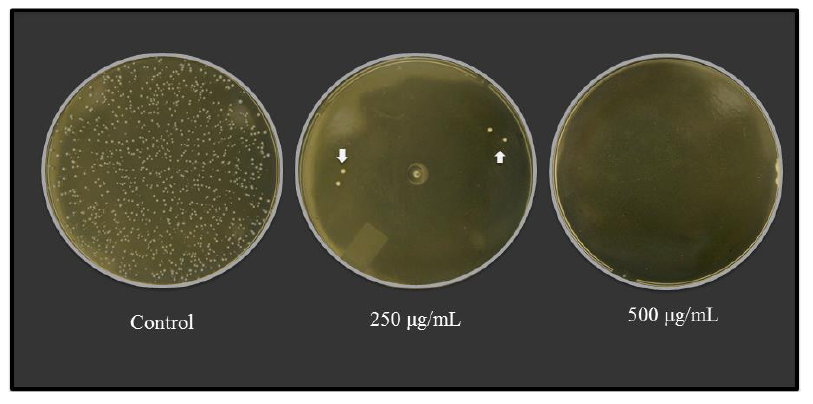
Agar BHI showing presence or absence of *S. mutans* CFU representing the MIC and MIB of zerumbone against *S. mutans*. Results are representative of three independent experiments performed in triplicate.Arrows: CFU.

**Supplementary figure 2.**
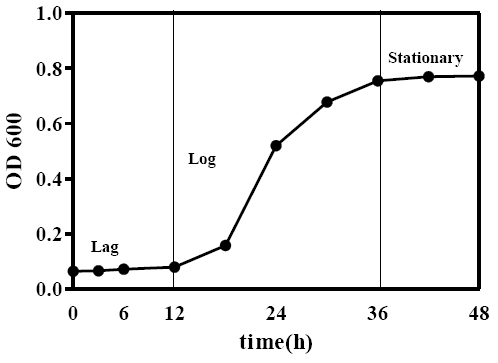
Bacterial growth curve of *S. mutans* cells gown in BHI broth for 48 hours at 37 in atmosphere of 5% CO_2_ at 37° C.

## Bibliography

1. Zakaria ZA, Yob NJ, Jofrry SM, Affandi MMRMM, Teh LK, Salleh MZ. Zingiber zerumbet (L.) Smith: A review of its ethnomedicinal, chemical, and pharmacological uses. Evidence-based Complement. Altern. Med. 2011.

2. Lang G, Buchbauer G. A review on recent research results (2008-2010) on essential oils as antimicrobials and antifungals. A review. Flavour Fragr. J. 2012. p. 13–39.

3. Lobo PLD, Fonteles CSR, Marques LARV, Jamacaru FVF, Fonseca SGDC, De Carvalho CBM, et al. The efficacy of three formulations of Lippia sidoides Cham. essential oil in the reduction of salivary Streptococcus mutans in children with caries: A randomized, double-blind, controlled study. Phytomedicine. 2014;21:1043–7.

4. Fakurazi S, Hairuszah I, Mohd Lip J, Shanthi G, Nanthini U, Shamima AR, et al. Hepatoprotective action of zerumbone against paracetamol induced hepatotoxicity. J. Med. Sci. 2009;9: 161–4.

5. Abdel Wahab SI, Abdul AB, Alzubairi AS, Mohamed Elhassan M, Mohan S. In vitro ultramorphological assessment of apoptosis induced by Zerumbone on (HeLa). J. Biomed. Biotechnol. 2009;2009.

6. Sulaiman MR, Tengku Mohamad TA, Shaik Mossadeq WM, Moin S, Yusof M, Mokhtar AF, et al. Antinociceptive activity of the essential oil of Zingiber zerumbet. Planta Med [Internet]. 2010;76: 107–12. Available from: http://www.ncbi.nlm.nih.gov/pubmed/19637111

7. M. N. Somchit. Zerumbone isolated from Zingiber zerumbet inhibits inflammation and pain in rats. J. Med. Plants Res. [Internet]. 2012;6: 753–7. Available from: http://www.academicjournals.org/JMPR

8. Sidahmed HMA, Hashim NM, Abdulla MA, Ali HM, Mohan S, Abdelwahab SI, et al. Antisecretory, gastroprotective, antioxidant and anti-helicobcter pylori activity of zerumbone from zingiber zerumbet (L.) smith. PLoS One. 2015;10.

9. Baby S, Dan M, Thaha ARM, Johnson AJ, Kurup R, Balakrishnapillaia P, et al. High content of zerumbone in volatile oils of Zingiber zerumbet from southern India and Malaysia. Flavour Fragr. J. 2009;24: 301–8.

10. Rahman HS, Rasedee A, Yeap SK, Othman HH, Chartrand MS, Namvar F, et al. Biomedical properties of a natural dietary plant metabolite, Zerumbone, in cancer therapy and chemoprevention trials. Biomed Res. Int. 2014.

11. Adbul ABH, Al-Zubairi AS, Tailan ND, Wahab SIA, Zain ZNM, Rusley S, et al. Anticancer activity of natural compound (Zerumbone) extracted from Zingiber zerumbet in human HeLa cervical cancer cells. Int. J. Pharmacol. 2008;4: 160–8.

12. Liu WY, Tzeng T-F, Liu I-M. Healing potential of zerumbone ointment on experimental full-thickness excision cutaneous wounds in rat. J. Tissue Viability [Internet]. Elsevier Ltd; 2017; 6–11. Available from: http://linkinghub.elsevier.com/retrieve/pii/S0965206X17300633

13. Petersen PE. Strengthening of Oral Health Systems: Oral Health through Primary Health Care. Med. Princ. Pract. [Internet]. 2014;23: 1–7. Available from: http://www.ncbi.nlm.nih.gov/pubmed/24525450

14. Bradshaw DJ, Lynch RJM. Diet and the microbial aetiology of dental caries: new paradigms. Int. Dent. J. [Internet]. 2013;63 Suppl 2: 64–72. Available from: http://www.ncbi.nlm.nih.gov/pubmed/24283286

15. Lynch DJ, Michalek SM, Zhu M, Drake D, Qian F, Banas JA. Cariogenicity of Streptococcus mutans Glucan-Binding Protein Deletion Mutants. Oral Health Dent. Manag. [Internet]. 2013;109: 191–9. Available from: http://www.omicsonline.com/open-access/cariogenicity-of-streptococcus-mutans-glucanbinding-protein-deletionmutants-2247–2452.1000512.pdf

16. Baehni PC, Takeuchi Y. Anti-plaque agents in the prevention of biofilm-associated oral diseases. Oral Dis. 2003;9 Suppl 1: 23–9.

17. Palombo EA. Traditional Medicinal Plant Extracts and Natural Products with Activity against Oral Bacteria: Potential Application in the Prevention and Treatment of Oral Diseases. Evid. Based. Complement. Alternat. Med. [Internet]. 2011;2011:680354. Available from: http://www.pubmedcentral.nih.gov/articlerender.fcgi?artid=3145422&tool=pmcentrez&rendertype=abstract

18. Ben El Hadj Ali I, Chaouachi M, Bahri R, Chaieb I, Boussaïd M, Harzallah-Skhiri F. Chemical composition and antioxidant, antibacterial, allelopathic and insecticidal activities of essential oil of Thymus algeriensis Boiss. et Reut. Ind. Crops Prod. 2015;77: 631–9.

19. Adams R. Identification of Essential Oil Components by Gas Chromatography / Mass Spectrometry. Identif. Essent. Oil Components by Gas Chromatogr. / Mass Spectrom. 1995;5.

20. Kumar SN, Lankalapalli RS, Kumar BSD. In vitro antibacterial screening of six proline-based cyclic dipeptides in combination with ??-lactam antibiotics against medically important bacteria. Appl. Biochem. Biotechnol. 2014;173: 116–28.

21. CLSI. M45-A2 Methods for Antimicrobial Dilution and Disk Susceptibility Testing of Infrequently Isolated or Fastidious Bacteria ; Approved Guideline — Second Edition. 2010.

22. Sartoratto A, Machado ALM, Delarmelina C, Figueira GM, Duarte MCT, Rehder VLG. Composition and antimicrobial activity of essential oils from aromatic plants used in Brazil. Brazilian J. Microbiol. 2004;35: 275–80.

23. Friedman PK, Kaufman LB, Karpas SL. Oral health disparity in older adults: Dental decay and tooth loss. Dent. Clin. North Am. 2014. p. 757–70.

24. Muts E-J, van Pelt H, Edelhoff D, Krejci I, Cune M. Tooth wear: a systematic review of treatment options. J. Prosthet. Dent. [Internet]. 2014;112: 752–9. Available from: http://www.ncbi.nlm.nih.gov/pubmed/24721500

25. Bagramian RA, Garcia-Godoy F, Volpe AR. The global increase in dental caries. A pending public health crisis. Am. J. Dent. 2009;22: 3–8.

26. Webster D, Taschereau P, Belland RJ, Sand C, Rennie RP. Antifungal activity of medicinal plant extracts; preliminary screening studies. [Internet]. J. Ethnopharmacol. 2008. p. 140–6. Available from: http://www.sciencedirect.com/science/journal/03788741

27. Kassebaum NJ, Bernabe E, Dahiya M, Bhandari B, Murray CJ, Marcenes W. Global burden of untreated caries: a systematic review and metaregression. J Dent Res [Internet]. 2015;94: 650–8. Available from: http://www.ncbi.nlm.nih.gov/pubmed/25740856%5Cn http://jdr.sagepub.com/content/94/5/650.full.pdf

28. Porto TS, Rangel R, Furtado NAJC, De Carvalho TC, Martins CHG, Veneziani RCS, et al. Pimarane-type diterpenes: Antimicrobial activity against oral pathogens. Molecules. 2009;14: 191–9.

29. Saleem M, Nazir M, Ali MS, Hussain H, Lee YS, Riaz N, et al. Antimicrobial natural products: an update on future antibiotic drug candidates. Nat. Prod. Rep. [Internet]. 2010;27: 238–54. Available from: http://dx.doi.org/10.1039/B916096E

30. da Silva JPC, de Castilho AL, Saraceni CHC, Díaz IEC, Paciencia MLB, Suffredini IB. Anti-Streptococcal activity of Brazilian Amazon Rain Forest plant extracts presents potential for preventive strategies against dental caries. J. Appl. Oral Sci. Rev. FOB. 2014;22: 91–7.

31. Freire ICM, P??rez ALAL, Cardoso AMR, Mariz BALA, Almeida LFD, Cavalcanti YW, et al. Atividade antibacteriana de ??leos essenciais sobre Streptococcus mutans e Staphylococcus aureus. Rev. Bras. Plantas Med. 2014;16: 372–7.

32. Takarada K, Kimizuka R, Takahashi N, Honma K, Okuda K, Kato T. A comparison of the antibacterial efficacies of essential oils against oral pathogens. Oral Microbiol. Immunol. 2004;19: 61–4.

33. Aguiar GP, Carvalho CE, Dias HJ, Reis EB, Martins MHG, Wakabayashi KAL, et al. Antimicrobial activity of selected essential oils against cariogenic bacteria. Nat. Prod. Res. [Internet]. 2013;27:1668–72. Available from: http://www.scopus.com/inward/record.url?eid=2-s2.0–84884285167&partnerID=tZOtx3y1

34. Santosh Kumar SC, Srinivas P, Negi PS, Bettadaiah BK. Antibacterial and antimutagenic activities of novel zerumbone analogues. Food Chem. 2013;141: 1097–103.

35. Hasan S, Danisuddin M, Khan AU. Inhibitory effect of zingiber officinale towards Streptococcus mutans virulence and caries development: in vitro and in vivo studies. BMC Microbiol. [Internet]. 2015;15:1. Available from: http://www.pubmedcentral.nih.gov/articlerender.fcgi?artid=4316655&tool=pmcentrez&rendertype=abstract

36. Jones RN, Anderegg TR, Deshpande LM. AZD2563, a new oxazolidinone: Bactericidal activity and synergy studies combined with gentamicin or vancomycin against staphylococci and streptococcal strains. Diagn. Microbiol. Infect. Dis. 2002;43: 87–90.

37. Anderl JN, Zahller J, Roe F, Stewart PS. Role of nutrient limitation and stationary-phase existence in Klebsiella pneumoniae biofilm resistance to ampicillin and ciprofloxacin. Antimicrob. Agents Chemother. 2003;47: 1251–6.

38. Gradelski E, Kolek B, Bonner D, Fung-Tomc J. Bactericidal mechanism of gatifloxacin compared with other quinolones. J. Antimicrob. Chemother. [Internet]. 2002;49:185–8. Available from: http://jac.oxfordjournals.org/content/49/1/185%5Cn http://jac.oxfordjournals.org/content/49/1/185.full.pdf%5Cn http://www.ncbi.nlm.nih.gov/pubmed/11751786

39. Nascimento MM, Lemos JA, Abranches J, Lin VK, Burne RA. Role of RelA of Streptococcus mutans in global control of gene expression. J. Bacteriol. 2008;190: 28–36.

40. Naidoo R, Patel M, Gulube Z, Fenyvesi I. Inhibitory activity of Dodonaea viscosa var. angustifolia extract against Streptococcus mutans and its biofilm. J. Ethnopharmacol. 2012;144: 171–4.

41. Silva F, Lourenço O, Queiroz J a, Domingues FC. Bacteriostatic versus bactericidal activity of ciprofloxacin in Escherichia coli assessed by flow cytometry using a novel far-red dye. J. Antibiot. (Tokyo). 2011;64: 321–5.

42. Waller SB, Madrid IM, Ferraz V, Picoli T, Cleff MB, de Faria RO, et al. Cytotoxicity and anti-Sporothrix brasiliensis activity of the Origanum majorana Linn. oil. Brazilian J. Microbiol. 2016;47: 896–901.

43. Lei J, Yu J, Yu H, Liao Z. Composition, cytotoxicity and antimicrobial activity of essential oil from Dictamnus dasycarpus. Food Chem. 2008;107: 1205–9.

44. Fallis A. Manual de toxicologia. J. Chem. Inf. Model. 2013.

